# Enhancement of synthetic mRNA translation efficiency through engineered poly(A) tails

**DOI:** 10.1101/2021.08.30.458298

**Authors:** Yusheng Liu, Hu Nie, Rongrong Sun, Jiaqiang Wang, Falong Lu

## Abstract

*In vitro* transcribed (IVT) mRNA represents a new class of drug in both therapeutics and vaccines. Improving the translation efficiency of IVT mRNA remains a core challenge for mRNA-based applications. Here, using IVT mRNAs with poly(A) tails containing non-A residues which were recently revealed to be widespread in RNA poly(A) tails^1,2^, we unexpectedly find that non-A residues can effectively promote the mRNA translation. To further support our finding, we provide evidence that non-A residues associated with enhanced mRNA translation efficiency transcriptome-wide in mouse and human cells. Together, our study provides a novel approach to enhance mRNA translation efficiency by inclusion of non-A residues in the mRNA poly(A) tails, holding great potential to promote mRNA-based therapeutics and vaccines.

## INTRODUCTION

In recent year, synthetic mRNA has become a promising approach for therapeutic gene delivery and vaccination^3,4^. Tremendous success has been achieved in mRNA vaccine against the ongoing global Covid-19 pandemic^5,6^. Great efforts have been invested to modify the mRNA to improve its stability and translation efficiency, including the 5’ cap, 5’UTR, 3’UTR, codon optimization, modified nucleosides and poly(A) tail^3^. The presence of a poly(A) tail is essential for mRNA stability, nuclear export and translation^7-16^. However, previous effort mainly focused on the length of the poly(A) tail, as previous knowledge of poly(A) tail mainly considers it as a structure component there and certain length ensures its translation and stability^3^.

Poly(A) tail information is lost in most transcriptome analysis because of the intrinsic difficulty of sequencing long homopolymeric stretches using second-generation sequencing platforms. Therefore, the roles of RNA poly(A) tails in regulating the fate and functions of RNAs are highly understudied. Researchers have used the TAIL-seq and PAL-seq (poly(A)-tail length profiling by sequencing) methods to estimate the lengths of poly(A) tails on the Illumina platform^17,18^. Studies of RNA poly(A) tails in oocytes of multiple species, including *Drosophila, Xenopus* and mice, revealed a clear link between the length of the poly(A) tail and the efficiency of mRNA translation^18-22^. Surprisingly, the length of a poly(A) tail, as long as it is longer than 20 nt, minimally affects mRNA translation in somatic cells^23^, leading to the question of whether poly(A) tails can regulate translation other than being a structural component of mRNA in somatic cells.

TAIL-seq revealed that the 3’ ends of mRNA poly(A) tails can be modified by end extension with other bases that can regulate the RNA stability, constituting another layer of mRNA stability regulation^17,24-28^. Short deadenylated poly(A) tails can be terminally U modified by TUT4/7, which can mark the target RNA for degradation^24^. In contrast, G modification catalyzed by TENT4A/B can protect the poly(A) tail from active deadenylation by the CCR4-NOT complex to stabilize the modified mRNA^25^.

The base composition within the body of poly(A) tails remained largely unknown until the recent development of the PAIso-seq and FLAM-seq methods on the PacBio platform, which allow long homopolymers to be sequenced accurately^1,2,29^. These methods revealed a novel group of RNA modifications, non-A residues in the RNA poly(A) tails (referred as non-A residues here after), that are widespread in the RNA poly(A) tails in mouse oocytes and human cell lines^1,2^, suggesting the existence a novel layer of RNA post-transcriptional regulation through non-A residues. Here, we reveal that non-A residues can enhance translation efficiency of IVT mRNA. We further provide evidence for this translation-enhancing function of non-A residues using unbiased genome-wide translation analysis. Together, we reveal a novel approach to enhance the translation of IVT mRNA using non-A residues, which will pave a new way of great potential to promote mRNA-based therapeutics and vaccines.

## RESULTS

### Non-A residues promote translation efficiency of IVT mRNA

Immediately after we and the Rajewsky lab identified non-A residues in mouse and human cells^1,2^, we asked what are the roles of these newly identified RNA residues in mRNA function. We synthesized mRNAs encoding fluorescent reporters with poly(A) tails of the same length but with different numbers of non-A residues using IVT (Fig. 1a). Then we transfected these IVT mRNAs into HeLa cells and monitored the encoded reporter protein and mRNA levels (Fig. 1b). RNA poly(A) tail is traditionally known as pure A sequence that can support translation. Therefore, our initial hypothesis was that mixed incorporation of nucleotides identified in the poly(A) tails might impair the role of poly(A) tail in supporting translation, leading to decreased or aborted translation. However, we unexpectedly observed obvious higher levels of EGFP fluorescence in cells transfected with EGFP mRNAs containing non-A residues compared with those transfected with EGFP mRNAs with a pure A tail (Fig. 1c), which was opposite to our initial hypothesis. Then, we used fluorescent-activated cell sorting (FACS) to quantify the level of florescent reporter translated from mRNA with non-A98-2 poly(A) tail and pure A tail. We found higher level of EGFP fluorescence for mRNA with the non-A98-2 tail; the same observation was seen when the mCherry reporter was used instead of EGFP (Fig. 1d). These results demonstrate that the non-A residues increase the protein production of multiple IVT mRNAs which is not limited to one special mRNA coding sequence.

**Figure 1.**
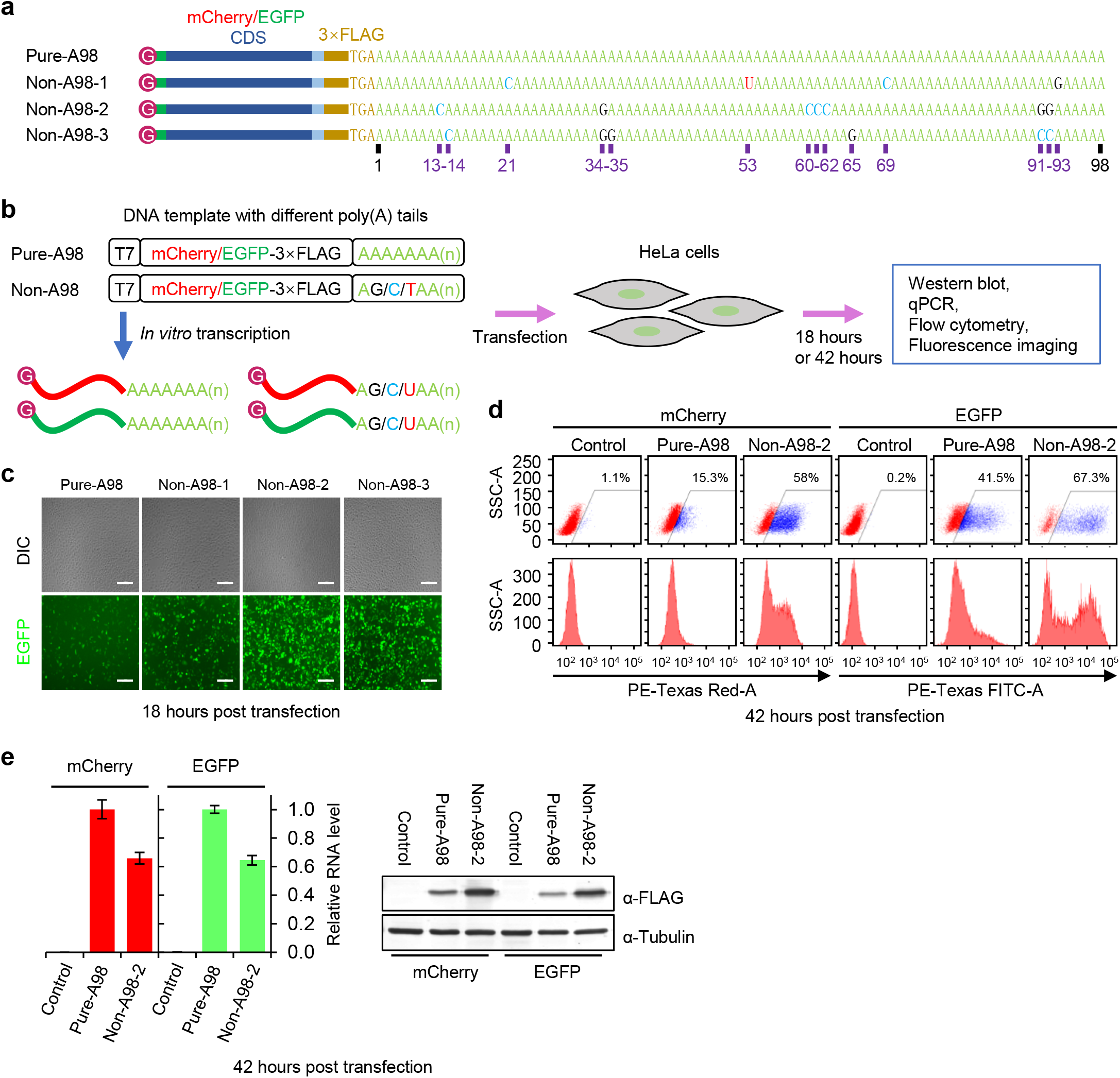
non-A residues promote translation of IVT mRNA. **a**, Schematic of reporter mRNAs with different non-A residues in poly(A) tails of the same length. **b**, Schematic of the translation efficiency reporter system based on transcripts with different types of poly(A) tails. **c**, Images of HeLa cells 18 hours post transfection with reporter mRNAs with different poly(A) tails. Scale bar, 100 μm. **d**, FACS quantification of reporter mRNA expression in HeLa cells 42 hours post transfection. **e**, Quantification of mRNA level by qRT-PCR (left) and protein level by western blot (right, anti-FLAG antibody was used to detect the FLAG tag for all reporters) 42 hours post transfection of HeLa cells with the reporter mRNA.

To investigate whether the increase of protein production by non-A residues is through increased translation efficiency or RNA stability, we then quantified the protein and mRNA levels within these cells by western blot and quantitative PCR (qPCR), respectively. Consistent with the observations under the microscope and FACS, the cells transfected with mRNAs containing non-A residues showed an increased amount of mCherry and EGFP protein production (Fig. 1e). qPCR analysis did not show higher mRNA levels for samples containing non-A residues, indicating that increased protein production was caused by increased translation efficiency rather than increased mRNA stability (Fig. 1e). To further confirm this effect of non-A residues in promoting translation, we also tested different time points after mRNA transfection and found consistent stronger fluorescence from 6 to 120 hours (Extended Data Fig. 1). Taken together, these data clearly demonstrate that non-A residues can greatly enhance the translation of synthetic mRNA.

### non-A residues are associated with high translation efficiency *in vivo*

The above findings showed enhanced translation efficiency by non-A residues in IVT mRNAs. In order to support the general function of non-A residues in enhancing translation that we unexpectedly discovered, we asked whether non-A residues presented in endogenous mRNAs can enhance the translation efficiency of these mRNAs. Therefore, we investigated the relationship between translation and non-A residues using the available ribosome engagement data and the non-A residues data for mouse GV oocytes and HeLa cells,^2,19,23,30^.

Consistent with previous observations^18,19,23,31^, poly(A) tail length was positively correlated with translation efficiency as measured by ribosome engagement in mouse GV oocytes (Fig. 2a), whereas no such correlation was observed in HeLa cells (Fig. 2b). This global positive association between poly(A) tail length and translation efficiency is seen only in reproductive cells but not in somatic cells^23^. In addition, global translation efficiency varies a lot during different stages of mitotic cell cycle, while the poly(A) tail length changed only moderately for a small proportion of genes^23^. However, we noticed extensive changes in non-A residues during somatic cell cycle^32^. Since poly(A) tail length does not contribute to translational regulation in somatic cells as seen in reproductive cells, we asked whether the non-A residues, the base composition of poly(A) tails, affect the translation efficiency.

**Figure 2.**
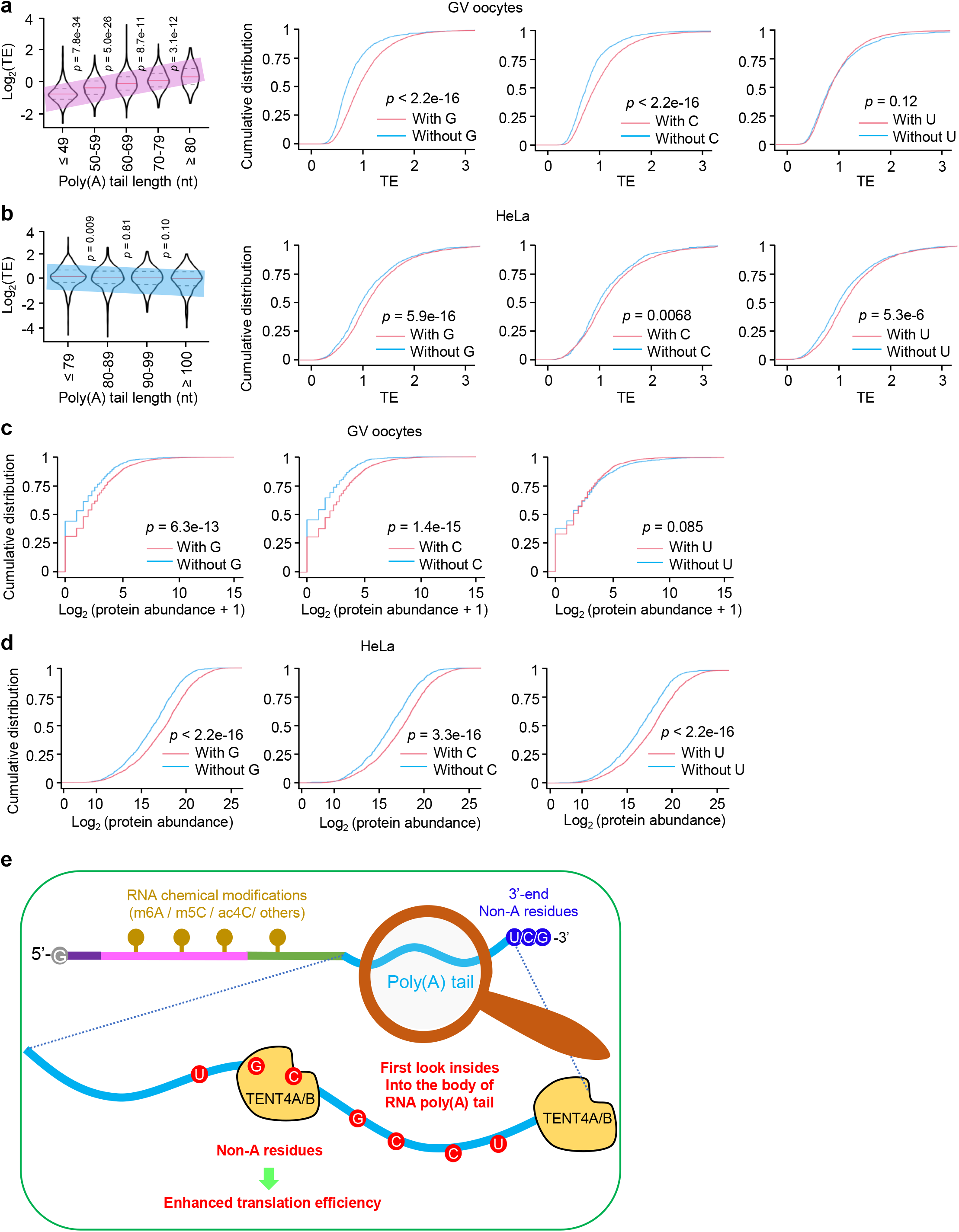
Endogenous non-A residues are associated with enhanced translation efficiency. **a, b**, Genome-wide relationship between poly(A) tail length^1,2^ and ribosome engagement (TE) (left) and cumulative distribution function (CDF) plots of mRNA-normalized TE for genes with or without non-A residues (right) in mouse GV oocytes^19^ (gene number: with G, n = 4,104; without G, n = 1,363; with C, n = 4,109; without C, n = 1,358; with U, n = 4,116; without U, n = 1,351) **(a)**, and HeLa cells^23^ (gene number: with G, n = 2,075; without G, n = 810; with C, n = 2,323; without C, n = 562; with U, n = 1,884; without U, n = 1,001) **(b)**. For GV oocyte sample, genes with 25% or more poly(A)+ mRNA containing G, C, or U residues were considered as genes with G, C, or U residues; while those containing less than 25% were considered as genes without G, C, or U residues. For HeLa sample, the overall level of non-A residues was much lower than that in GV oocytes. Therefore, genes with or without poly(A)+ mRNA containing G, C, or U residues were considered as genes with or without G, C, or U residues. Genes with at least 10 ccs reads were included in the analysis. In the violin plots, median values are represented by red lines and the first and third quartiles are represented by black, dashed lines. *p* value was calculated by two-tailed Student’s *t* test for the relationship between length and TE and by Kolmogorov-Smirnov (KS) test for the relationship between non-A residues and TE. **c, d**, CDF plots of protein abundance quantified by mass spectrometry for genes with or without non-A residues in GV oocytes^33^ (gene number: with G, n = 2,307; without G, n = 768; with C, n = 2,308; without C, n = 767; with U, n = 2,306; without U, n = 769) **(c)**, and HeLa cells^34^ (gene number: with G, n = 2,398; without G, n = 1,029; with C, n = 2,694; without C, n = 733; with U, n = 2,166; without U, n = 1,261) **(d)**. The *p* value was calculated by KS test. **e**, A model for the full view of a complete single mRNA molecule. The magnified part highlights the non-A residues in RNA poly(A) tails, which are conserved in eukaryotes. Other known RNA modifications such as m6A, m5C, ac4C and terminal non-A modifications are also labeled. The non-A residues can enhance the translation of the mRNA.

Interestingly, genes with G and C residues showed significantly higher translation efficiencies in both germ cells and somatic cells (Fig. 2a, b). However, U residues were not always associated with increased translation efficiency, especially in mouse GV oocytes (Fig. 2a, b). This may be related to the much higher abundance of U residues in GV oocytes, especially consecutive oligo-U residue (Extended Data Fig. 2), which might result in uridylation mark RNA for degradation^24,27^. Given that mRNA translation ultimately results in the accumulation of proteins, we investigated the relationship between non-A residues and protein abundance in mouse GV oocytes and HeLa cells, for which the mass-spectrometry based proteomic data were available^33,34^. The results showed that proteins encoded by genes with non-A residues showed significantly increased abundance in both GV oocytes and HeLa cells, which is consistent with the ribosome engagement data (Fig. 2c, d). These results reveal that the non-A residues are indeed prevalent regulators of translation of endogenous mRNAs in both germline and somatic cells, demonstrating that poly(A) tails can be critical regulators of mRNA translation rather than being simple structural components. Therefore, these unbiased genome-wide analysis of endogenous mRNA non-A residues and translation well support our finding that non-A residues promote translation of IVT mRNA.

## DISCUSSION

In recent years, mRNA-based therapeutics have become a new focus in the pharmaceutical industry^3,4^. Compared with protein-based therapy, mRNA-based therapy is much more cost-effective because mRNAs are easy to design and produce. mRNA vaccines allow the fastest pre-clinical development, and have proven great potential in responding to emerging outbreaks of infectious diseases^5,6,35,36^. For example, mRNA vaccines designed and synthesized within one month from Moderna and BioNTech were among the first vaccines progressing into phase 1/2 clinical trials during the current ongoing global Covid-19 pandemic^37-40^. Moreover, these mRNA vaccines are the first approved vaccines and proven of exceptional efficacy in preventing Covid-19^5,6^. Improving translation efficiency has been one of the main technical challenges in mRNA-based therapeutics and vaccines^3,41^. In this study, we demonstrate that non-A residues are conserved and abundant novel RNA modifications in diverse eukaryotes that can enhance mRNA translation (Fig. 2e). The discovery that non-A residues enhance translation of IVT mRNA provides a completely new way to promote the development of mRNA-based therapeutics and vaccines.

In order to further confirm our finding that non-A residues enhance translation of IVT mRNAs, we provide evidence that non-A residues associate with enhanced mRNA translation in human and mouse cells, revealing the non-A residues as endogenous mRNA translation regulating factors. Our recent study reveal the conservation and prevalence of non-A residues in mRNA poly(A) tails across eukaryotes, suggesting universal functions of them in regulating diverse biological processes^29^. As the above data reveal that non-A residues enhance mRNA translation in human and mouse cells, the prevalence of non-A residues suggest that non-A residues may share similar function in enhancing mRNA translation across eukaryotic species. Therefore, the including of non-A residues in the synthetic mRNAs will likely be a novel approach to enhance the protein production in mRNA-based applications across the eukaryotic species. What enzymes are responsible for the incorporation of these non-A residues is an interesting question to answer for mechanistic study *in vivo*. The mechanisms by which non-A residues promote mRNA translation and whether G, C and U residues are using different mechanisms are the next important questions awaiting further investigations.

Previous methods to improve IVT mRNA stability, translational properties and immunogenicity largely relies on chemically modified nucleosides in mRNA, such as pseudouridine, 1-methyl-pseudouridine 2-thiouridine, 5-methyluridine, 5-methylcytidine or N6-methyladenosine^3,4^. However, chemically modified nucleosides such as pseudouridine and N6-methyladenosine severely decrease synthesis efficiency and increase error rate associated during IVT mRNA synthesis with the use of chemically modified nucleosides^42^. Here, we discover a completely new way to improve mRNA translation by mixed incorporation of non-A residues within the mRNA poly(A) tails. These non-A residues are regular nucleotides, which can be easily designed into the mRNA synthesis template to ensure 100% reproducible incorporation into the final full length mRNA products. Moreover, as this new approach is different from all the current available approaches in improving the pharmacokinetics of mRNA drugs, it can be readily incorporated into every existing mRNA developing pipelines together with other methods, such as combined with the use of modified nucleosides that reduce immunogenicity and increase stability. In summary, the new approach described in the current study using non-A residues to enhance IVT mRNA translation will have considerable translational potential in pushing forward the mRNA therapeutics and vaccines.

## METHODS

### mRNA preparation, cell culture and transfection

The template used for *in vitro* transcription (IVT) were amplified by PCR from the pcDNA3.1-EGFP/mCherry-3XFlag plasmids using T7-F forward primer and different reverse primers (Supplementary Table 1). The capped mRNAs were *in vitro* synthesized with HiScribe T7 ARCA mRNA Kit (NEB) and purified with an RNA Clean & Concentrator-25 kit (Zymo Research) according to the manufacturers’ instructions. The purified mRNA was dissolved in RNase-free water and stored at -80 °C until use. HeLa cells were grown in DMEM (Invitrogen) containing 10% FBS (Gibco) at 37 °C in a humidified 5% CO_2_ incubator. For mRNA transfection, HeLa cells were transfected with Lipofectamine MessengerMAX reagent (Invitrogen). The transfected cells were used for imaging and FACS analysis or harvested for other downstream experiments at the indicated time points.

### Western blot analysis

HeLa cells were lysed with SDS loading buffer containing protease inhibitor cocktail (Roche), boiled and separated by SDS-PAGE. The samples were then transferred onto PVDF membrane. The membrane was incubated with primary antibody (mouse anti-FLAG antibody (Sigma, F1804; 1:1000); mouse anti-Tubulin (Sigma, T9026; 1:1000) overnight at 4 °C. The next day, after washing 3 times with TBST, the membrane was incubated with fluorescent dye-labeled secondary antibody (LI-COR, 925-32210, 1:1000) for 1 hour at RT. The membrane was then scanned using an Odyssey infrared scanner (LI-COR).

### Quantitative real-time RT-PCR

The total RNA was isolated from HeLa cells with Direct-zol RNA MicroPrep (Zymo Research) according to the manufacturer’s instructions. The SuperScript II Reverse Transcriptase (Invitrogen) was used to synthesize cDNA from RNA. Quantitative PCR was performed using TB Green Premix Ex Taq (Takara) with specific primers (Supplementary Table 1). Measurements were performed with three independent biological replicates.

### Detection of non-A residues in poly(A) tails

To minimize errors introduced by sequencer, we used clean CCS reads with at least ten passes to call non-A residues in poly(A) tails. G, C, U (presented as T in CCS reads) were counted in poly(A) tail of each CCS reads. The percentage of transcripts with non-A residues of a gene was the number of CCS reads with at least one G, C or U residue divided by the total number of CCS reads with poly(A) tails of length at least 1 nt for the given gene.

### TE and the protein abundance analysis

Translation efficiency (TE) was calculated based on ribosome engagement data and related RNA-seq data (GSE135525 for mouse GV oocyte RiboTag/RNA-seq^19^, GSE79664 for HeLa Ribo-seq^23^). Genes with at least ten reads measured in both ribosome engagement and RNA-seq data were included in the analysis. CPM (counts per million) were calculated to represent the relative gene expression levels. Translation efficiency was calculated by dividing the CPM of ribosome engagement fragments to that of the total mRNA. Protein copy-number estimates assigned to each gene were used to represent the protein abundance in HeLa cells^34^. The number of observed spectra assigned to each gene were used to represent the protein abundance in mouse GV oocytes^33^. Other than the PAIso-seq data obtained in this study, HeLa cell poly(A) tail information from the previous poly(A) tail analysis were used for the translation efficiency and protein abundance analysis^1^.

## Data availability

The ccs data in bam format from PAIso-seq1 experiments (GV oocytes) will be available at Genome Sequence Archive hosted by National Genomic Data Center. This study also includes analysis of the following published data: Legnini et al. (Gene Expression Omnibus database (GEO) accession no. GSE126465), Liu et al. (Sequence Read Archive database (SRA) accession no. PRJNA529588), Luong et al. (GEO accession no. GSE135525), Park et al. (GEO accession no. GSE79664), and Bekker-Jensen et al. (ProteomeXchange Consortium no. PXD004452). Custom scripts used for data analysis will be available upon request.

## Acknowledgements

This work was supported by the Strategic Priority Research Program of the Chinese Academy of Sciences (XDA24020203), the National Key Research and Development Program of China (2018YFA0107001), National Natural Science Foundation of China (31970588), Natural Science Foundation of Heilongjiang province (YQ2020C003), the China Postdoctoral Science Foundation (2020M670516, 2020T130687), and the State Key Laboratory of Molecular Developmental Biology.

## Author Contributions

Yusheng Liu, Jiaqiang Wang and Falong Lu conceived the project and designed the study. Yusheng Liu and Rongrong Sun performed the synthetic mRNA study in culture cells. Yusheng Liu, Hu Nie, Jiaqiang Wang and Falong Lu analyzed the sequencing data. Yusheng Liu and Jiaqiang Wang organized all figures. Yusheng Liu, Jiaqiang Wang and Falong Lu supervised the project. Yusheng Liu, Jiaqiang Wang and Falong Lu wrote the manuscript with the input from the other authors.

## Competing interests

Y.L. and F.L. are named inventors on a patent application filed by Institute of Genetics and Developmental Biology covering the discovery in this manuscript.

## FIGURE LEGENDS

**Extended Data Fig. 1.**
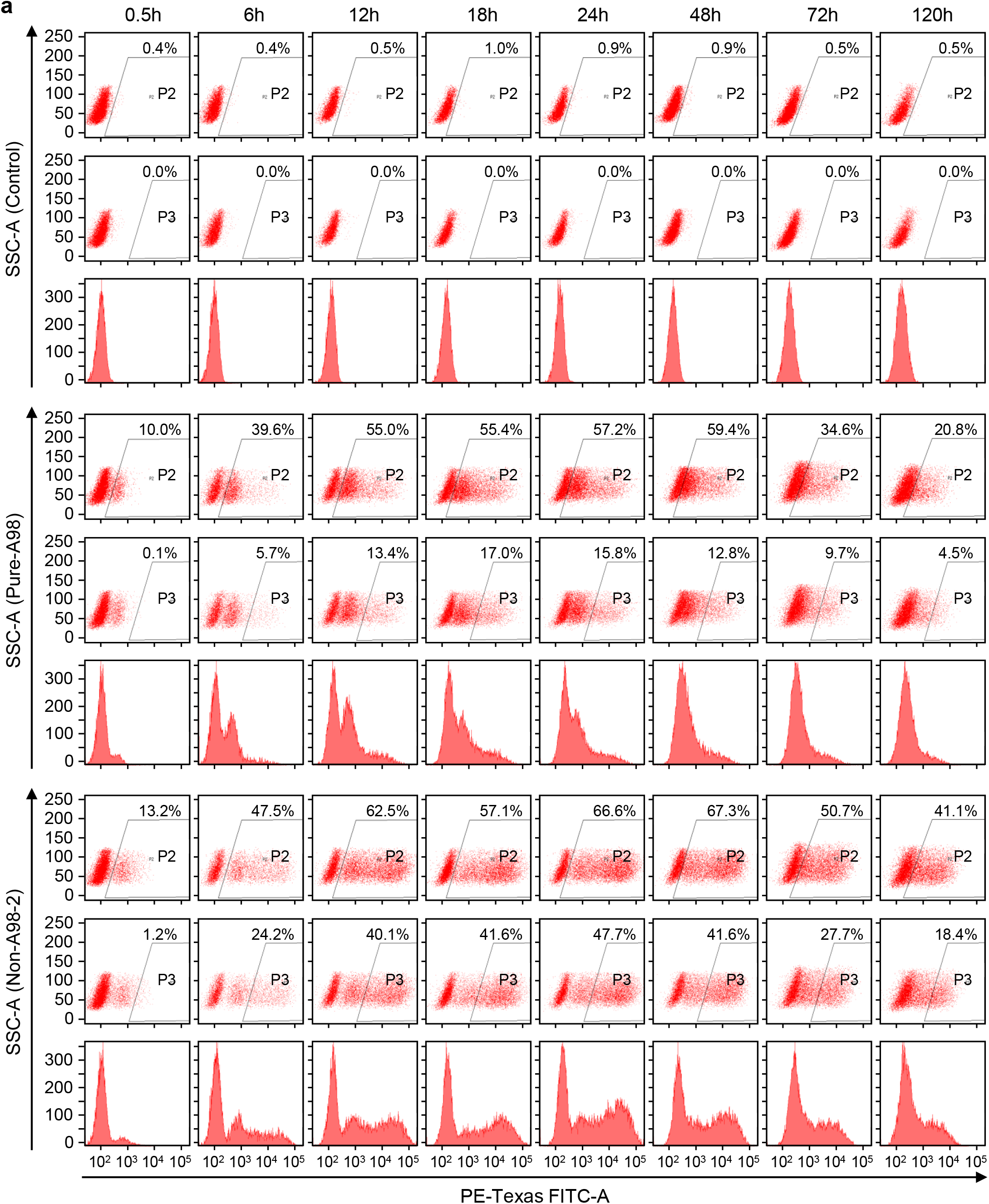
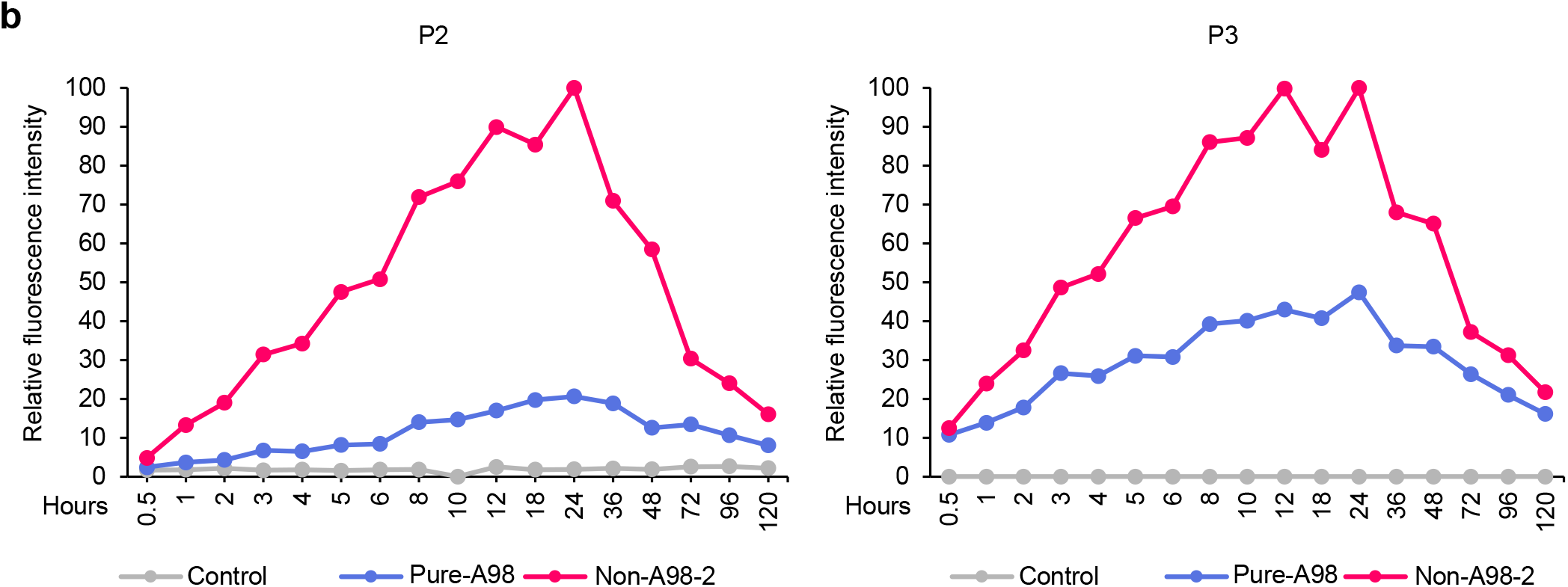
Quantification of fluorescent reporter translated from IVT mRNA at different time points. **a**, Quantification of reporter EGFP mRNA expression in HeLa cells by FACS at different time points post transfection. Control: cells not transfected with reporter mRNA. PURE-A98 and Non-A98-2: cells transfected with reporter mRNAs with the PURE-A98 and Non-A98-2 poly(A) tails, respectively, as shown in Fig. 1a. **b**, Relative fluorescence intensity quantification of reporter mRNA expression in HeLa cells at different time points by FACS shown in panel a.

**Extended Data Fig. 2.**
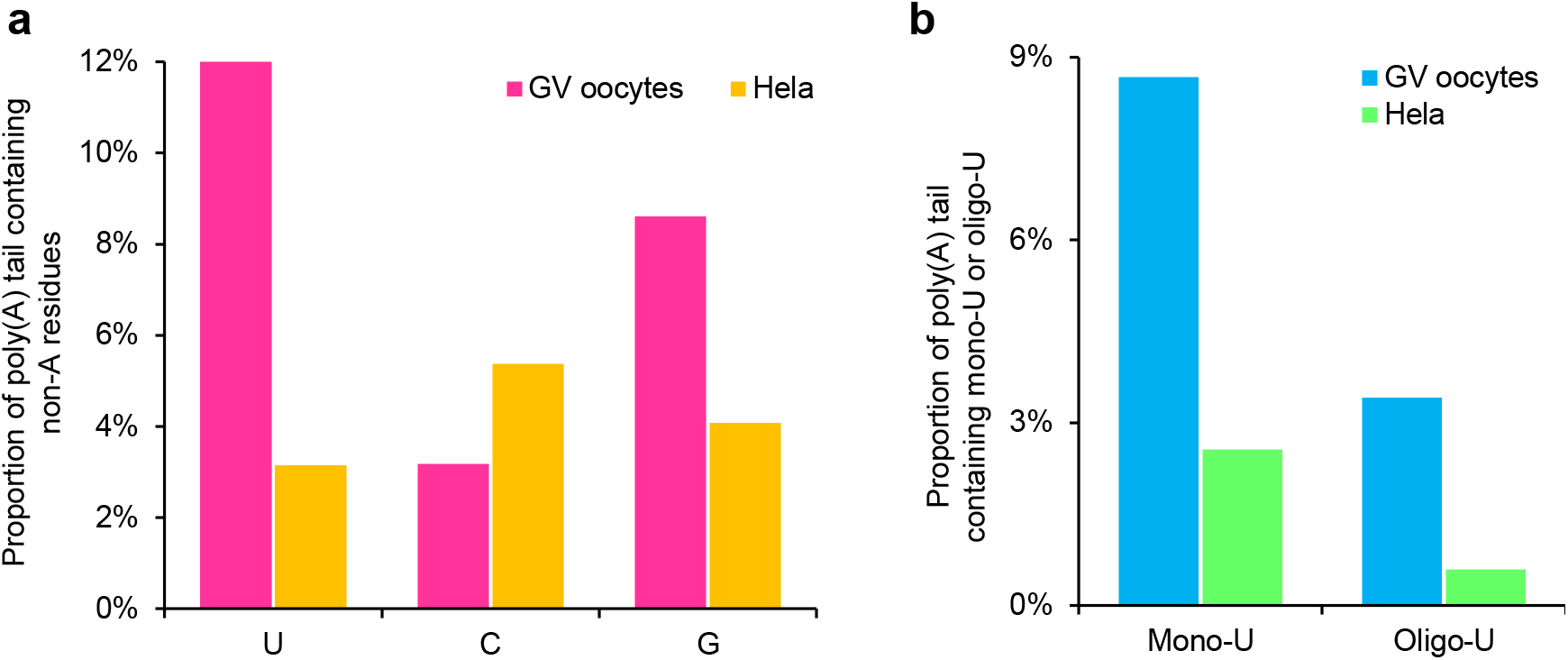
Abundance of non-A residues. Proportion of the transcripts containing non-A residues **(a)**, and mono- and oligo-Us **(b)** in mouse GV oocytes^2^ and HeLa cells^1^.

## Notes

### Competing Interest Statement

The authors have declared no competing interest.

